# A 3.8 Å Resolution Cryo-EM Structure of a Small Protein Bound to a Modular Imaging Scaffold

**DOI:** 10.1101/505792

**Authors:** Yuxi Liu, Duc Huynh, Todd O. Yeates

**Affiliations:** UCLA Department of Chemistry and Biochemistry; UCLA-DOE Institute for Genomics and Proteomics; UCLA Molecular Biology Institute; California NanoSystems Institute, UCLA

## Abstract

Proteins smaller than about 50 kDa are currently too small to be imaged by cryo-electron microscopy (cryo-EM), leaving most protein molecules in the cell beyond the reach of this powerful structural technique. Here we use a designed protein scaffold to bind and symmetrically display 12 copies of a small 26 kDa protein. We show that the bound cargo protein is held rigidly enough to visualize it at a resolution of 3.8 Å by cryo-EM, where basic structural features of the protein are visible. The designed scaffold is modular and can be modified through modest changes in its amino acid sequence to bind and display diverse proteins for imaging, thus providing a general method to break through the lower size limitation in cryo-EM.

## Main Text

Methods for visualizing macromolecules in atomic detail have transformed our understanding of molecular biology. Yet the leading techniques – X-ray crystallography, NMR, and electron microscopy – all face obstacles that limit their universal application. X-ray crystallography presents difficulties in crystallization, while NMR methods become challenging for very large macromolecules. For cryo-EM, despite recent technological advances that have revolutionized the field (reviewed in 1, 2), a lower size limitation has prevented its application to proteins smaller than about 50 kDa, which is larger than the average cellular protein. Overcoming this lower size barrier could bring electron microscopy close to the ultimate goal of a universally applicable method for structural biology.

The goal of visualizing small proteins by cryo-EM has motivated research efforts along multiple directions. The use of phase plates (3) enhances image contrast in cryo-EM and has enabled recent reports that pushed the lower size limit for imaging. Two recent structural studies on human hemoglobin (64kDa) and streptavidin (52kDa), both at a resolution of 3.2 Å (4, 5), have begun to approach the 38 kDa theoretical limit proposed in 1995 (6). An alternative approach is to design molecular ‘scaffolding’ systems – i.e. molecules of known structure that are large enough to visualize in atomic detail by cryo-EM, and simultaneously able to bind and display a smaller protein molecule of interest, effectively making the smaller ‘cargo’ protein part of a larger assembly that can be structurally elucidated as a whole (7–11). Two key challenges have hindered the development of useful cryo-EM scaffolds: rigidity and modularity. The cargo protein must be bound and displayed rigidly so that it does not become smeared out during reconstruction of the full structure. And to be practical the scaffold must be able to bind diverse cargo proteins with minimal re-engineering efforts.

In a recent work, we developed a scaffold that addresses the requirements for rigid display and modularity, while also exploiting the advantage of high symmetry to mitigate the common problem of preferred particle orientation in cryo-EM (10). Our scaffolding system uses a DARPin (Designed Ankyrin Repeat Protein) as an ‘adaptor’ component that is genetically fused by way of a continuous alpha helical connection to a central core comprised of a designed symmetric protein cage with cubic symmetry (12 orientations in symmetry T, Fig. 1) (12). In prior work, DARPins have been developed as a facile framework for sequence diversification and selection, by phage display or other laboratory evolution methods, for binding to a wide range of target proteins (13–20). With these ideas put together, our scaffold presents multiple symmetrically disposed copies of an adaptor protein whose sequence can be mutated to bind diverse cargo proteins for imaging. In the first structural study on this scaffolding system, referred to as DARP14, we analyzed the scaffold by itself without cargo bound and demonstrated that the DARPin adaptor component could be visualized by cryo-EM at a resolution ranging from 3.5 to 5.5Å (10). Here we present the first cryo-EM study where this designed scaffold is used to bind a cargo protein for imaging.

**Figure 1.**
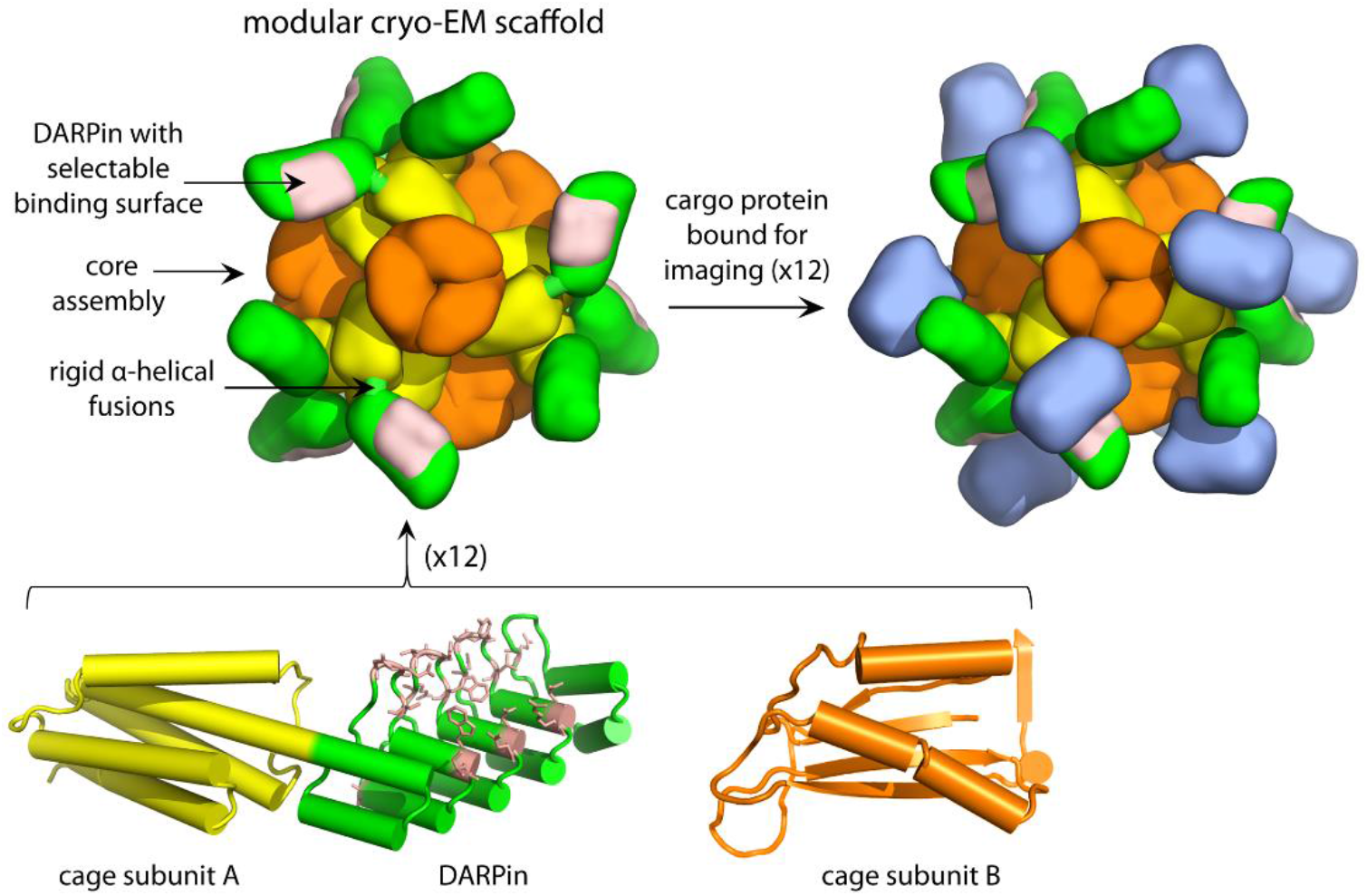
A scheme illustrating the design of the modular scaffold and its binding to target cargo proteins for cryo-EM imaging. The designed core assembly is composed of 12 copies of two protein subunits, A (yellow) and B (orange), in a tetrahedrally symmetric arrangement. Subunit A is genetically fused by a continuous alpha helical linker to a DARPin (green). Amino acid mutations (based on *in vitro* selection experiments) are inserted into the DARPin binding surface (pink) to confer tight binding of a cargo protein (blue) for cryo-EM imaging.

As a first test case we chose the 26 kDa green fluorescent protein (GFP) as a cargo molecule. GFP has been well-studied and DARPin sequences that bind various forms of GFP have been published (18,19). Our studies employed the superfolder variant (sfGFP) (21) with a V206A mutation. Compared to the prior study, we modified the DARP14 scaffold as follows. Motivated by the finding that the terminal repeats of the DARPin were more flexible and showed a compromised resolution, we adopted a set of mutations previously characterized to stabilize this region of the DARPin (22). Finally, to convert the scaffold to a form that would bind GFP – and thus illustrating the modularity of the system – we exchanged the amino acid sequences in the binding loops on the original DARP14 scaffold with a sequence known as 3G124 (18, 19). Taken together, these modifications make a symmetric scaffold that is specific to GFP. After binding GFP to the scaffold, the complex was purified and found by quantitative amino acid analysis to have nearly complete saturation of the binding sites on the scaffold; i.e. nearly 12 copies of GFP on each cubically symmetric scaffold. Negative stain EM showed assembled particles of the expected size and symmetry (Fig. S1A).

To analyze the three-dimensional structure, we collected 1929 movie images of this complex under cryogenic conditions using a Titan Krios (Fig. S1B). After initial data processing, 81,319 particles were selected for 3D analysis based on their 2D class averages (Fig. S2). The 2D class averages showed strong density for the symmetric core of the scaffold. Moreover, there was clear additional density for the DARPin adaptor and the bound GFP (Fig. 2A, compare to Fig. 2B in (10)). Some flexibility was evident for the DARPin and the GFP cargo. The density in these regions was generally weaker and more diffuse, indicative of multiple conformations and calling for special attention in the later analysis.

**Figure 2.**
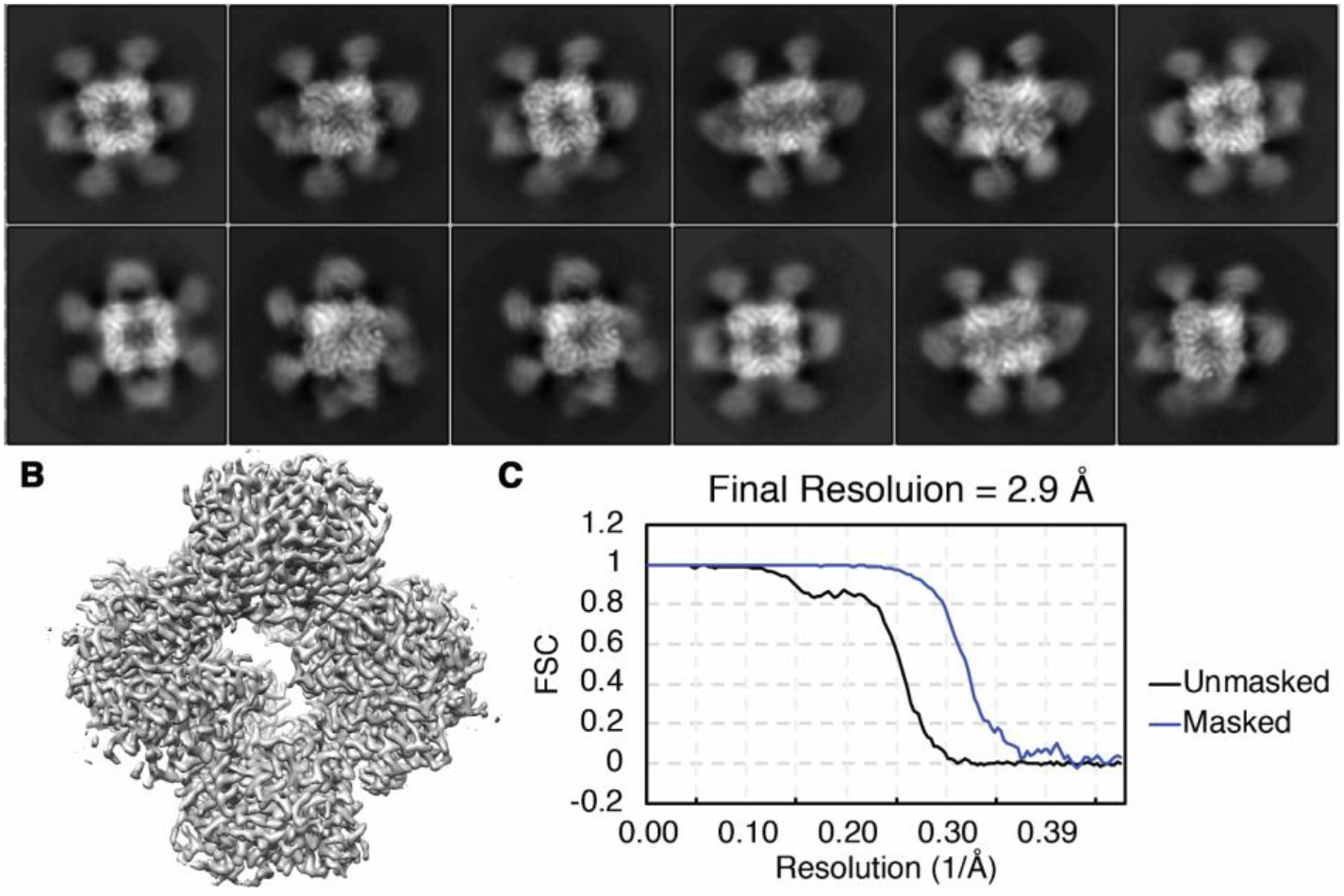
Cryo-EM data on GFP bound scaffolds. (A) 2D class images of the particles show strong features for the tetrahedral core along with clear but more diffuse density for the DARPin and the bound GFP cargo. The 3-D density reconstructed only around the core structure (B) and the corresponding gold-standard Fourier shell correlation (FSC) curve (C) are shown.

An initial 3D image reconstruction was performed using the CryoSPARC program (23), focusing on the symmetric core of the scaffold with enforced T symmetry (Fig. 2B, S2). This map exhibited an overall resolution of 2.9 Å with good amino acid side chain features all around (Fig. 2B, C). Refining the structure of the scaffold core against this map showed that the core structure is largely undisturbed relative to our previous cryo-EM model, with an overall RMSD of 0.90 Å between the two (10). To reconstruct the attached DARPin and GFP components, the particles were then passed to the RELION program (24, 25) to perform signal subtraction and masked classification as previously described (26, 27). After expanding the particles by T symmetry and subtracting away the density corresponding to 11 of the 12 DARPin adaptors and their bound GFP molecules, the remaining density was classified again in 3D without alignment. Interestingly, about 13.3% of the particles have very weak densities for the adaptor and cargo, likely indicating that these instances suffered from very different (i.e. bent) orientations for the protruding components. Three classes had good density for the DARPin and the bound GFP components, with slight differences in orientation for the distinct classes. They showed density around the previously observed secondary contact point between the DARPin and the scaffold core (Gly 187 on DARPin and Gly 108-Thr 109 on subunit A of the scaffold core, Fig. S3). We performed two sequential masked refinements on particles from class 1, followed by a multibody refinement. The masks used in the refinements increasingly focused on a single DARPin and its bound GFP (Fig. S2). And the multibody refinement searched the DARPin and GFP orientation locally while being constrained by its position relative to the symmetric core. The final reconstruction reached 3.5 Å in overall resolution, with the scaffold core at a somewhat higher resolution than the DARPin or GFP (Fig. 3A-C).

**Figure 3.**
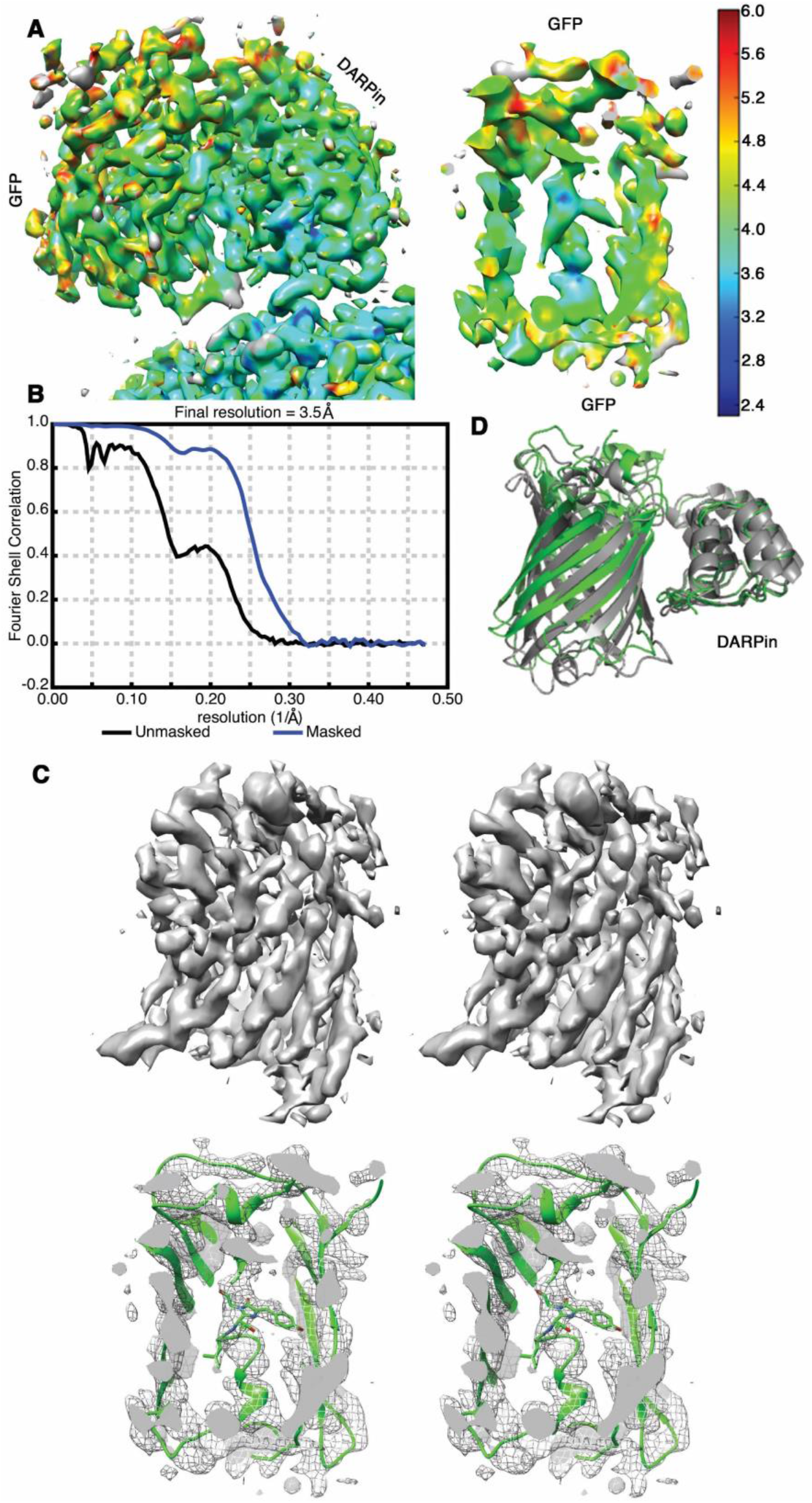
Near-atomic resolution maps of the DARPin adaptor and its bound cargo protein, GFP. (A) A density map obtained following multi-body refinement in RELION is shown colored by local resolution in a side view (left) and a sliced view (right), showing the central helix in the center of the GFP beta barrel. (B) Gold-standard FSC curve for the reconstruction. (C) Two wall-eyed stereo views of the sharpened GFP density map. The full GFP density is shown at the top; a sliced view with a fitted GFP atomic model is shown at the bottom. The fluorophore and Leu 64 (shown in sticks) in the central helical region have clear surrounding density. (D) Comparison between the DARPin plus GFP model in this study (green) and the previously reported 3G124 eGFP crystal structure (grey, PDB 5MA8). The two structures are aligned on the DARPin portion.

Despite the presence of some flexibility in the DARPin and bound cargo (GFP), the resolution that was achievable for these components was remarkably good. The local resolution for the DARPin ranges from 3.3 Å to 3.8 Å (1^st^ to 3^rd^ quartile values) (28). This represents a notable improvement over our initial report on the scaffold by itself, likely due to the stabilizing mutations introduced into the terminal repeat of the DARPin and the presumed stabilizing effects of binding to the cargo protein, GFP. The local resolution for the bound GFP ranges from 3.6 Å to 5.1 Å (1^st^ to 3^rd^ quartile values) with a median value of 3.8 Å. Key structural features of the protein are clear. The side of GFP contacting the DARPin shows a better resolution than the outward-facing side. In the regions of better resolution, the slanting beta strands in the GFP beta barrel are separate and clearly traceable, and side chain densities for some of the bulky residues are also visible. The alpha helical segment passing through the center of the β-barrel exhibits a local resolution of 3.5 Å, with side chain and fluorophore features clearly visible in the sharpened density map (Fig. 3C). This level of detail represents a new benchmark for imaging the structures of scaffolded small proteins by cryo-EM.

We undertook calculations to evaluate how much structural information could be extracted at the current resolution of 3.8 Å. It was possible to refine the DARPin with secondary structure restraints based on a reported unbound structure (PDB 6C9K) and to refine GFP as a rigid body based on a known crystal structure (PDB 4W6B). Refinement revealed that the individual ankyrin repeats of the DARPin adaptor expanded slightly further away from each other on the opposite side of the bound GFP. In addition, our bound superfolder GFP is slightly rotated compared to the eGFP bound to a DARPin with the same loop sequence (PDB 5MA8, Fig. 3D). Side chain densities of bulky residues at the GFP-DARPin interface are well-resolved. Overall, the final model shows a good fit to the electron density, with clear chain tracing in much of the structure, and agreement with side chain density in the higher resolution regions (Fig. 3C).

Our results illustrate a route for overcoming the critical lower protein size limit in cryo-EM. While the current resolution allows for visualization of useful structural features of a protein, it falls just short of providing the atomic detail required to elucidate the structures of novel proteins. That goal will likely be possible by improving the resolution by about 0.5 to 1 Å, to reach the 3 Å range. Our scaffolding system offers strategies for realizing such an incremental improvement. In particular, the high symmetry of the system admits additional design components for introducing stabilizing connections between symmetry-related pairs of the protruding DARPin components. Only modest advances will be needed in order to visualize atomic detail for small proteins. The value of the DARPin system is also demonstrated by another recent report of an engineered scaffolding system based on the D2 tetrameric aldolase utilizing a DARPin as the adaptor to image eGFP, albeit at a lower resolution (5-8 Å) (11). Among the varied strategies under development for overcoming the lower size limit in cryo-EM, scaffolding approaches are likely to offer advantages beyond simply increasing the effective size of the imaging target. Highly symmetric systems as developed here help mitigate the frequently-encountered problem of preferred particle orientation; even when a symmetric particle exhibits an uneven orientation distribution, its symmetrically repeating components are effectively viewed from multiple directions simultaneously. The presence of multiple copies of a cargo protein on each particle also assures that some will be away from the air-water interface, where structural integrity is often compromised due to poorly understood surface chemistry and mechanics (29, 30). Scaffold approaches also provide advantages in streamlined cryo-EM data processing, as a large symmetric core greatly simplifies the critical problem of establishing accurate particle orientations. Besides seeking improvements in resolution, future experiments that include more diverse targets will be important in order to demonstrate the versatility and imaging advantages conferred by cryo-EM scaffolding systems of the type described here.

## Ackdnowledgments

The authors thank Prof. Hong Zhou, Yanxiang Cui and Kang Zhou for assistance in cryo-EM data collection. We also thank David Boyer, Peng Ge, and Duilio Cascio for advice on cryo-EM data processing. We also acknowledge Andreas Plückthun for advice on DARPins, and Christopher Phillips, Taiana Maia De Oliveira, and Alexis Rohou for general discussion and advice on cryo-EM data collection and DARPins, and Mark Yeager and James Hogle for discussions on scaffolds.

## Author contributions

Y.L. and T.O.Y. designed research; Y.L. and D.H. performed the experiments; Y.L. and T.O.Y. analyzed the data; T.O.Y. supervised the research; Y.L. and T.O.Y. wrote the paper and D.H. provided revisions.

## Data and materials availability

Model coordinates and density maps will be released upon publication.

## MATERIALS AND METHODS

### Cloning and Protein expression

The sequence of the scaffold used in this paper, DARP14-3G124Mut5 (see SI text), was designed by replacing the original DARPin sequence (10) with the anti-GFP DARPin sequence, referred to as 3G124 (19), while adding the Mut5 mutations into the C-terminal repeat (22). The mutations are A299P, I3022L, S303A, L311I, I314V, L218A, N319A (as numbered in the DARP14 subunit A sequence). Superfolder GFP V206A (sfGFP V206A) was cloned with quick-change PCR to introduce the V206A mutation on a pET-22b vector carrying the superfolder GFP gene (21). DARP14-3G124Mut5 and sfGFP V206A were expressed separately in *E. coli* BL21 (DE3) cells in Terrific Broth at 20°C overnight upon 1mM IPTG induction at O.D. 0.6.

Upon collection of the cells, DARP14-3G124Mut5 and sfGFP V206A pellets were mixed at a 3:1 mass ratio, resuspended in resuspension buffer (50mM Tris, 250mM NaCl, 5mM imidazole, 5% (v/v) glycerol, pH 8.0) supplemented with DNAse, lysozyme, and protease inhibitor cocktail (Thermo Fisher Scientific) and lysed together by sonication. Cell lysate was first cleared by centrifugation at 20,000 x g for 20min and loaded onto a HisTrap column (GE Healthcare) pre-equilibrated with the same resuspension buffer. DARP14-3G124Mut5 bound with sfGFP V206A was eluted with a linear gradient to 500mM imidazole. Upon elution, 5mM DTT was added immediately. The eluted protein was concentrated and further purified by passing through a Superose 6 Increase 10/300 GL column (GE Healthcare), eluted with 10mM Tris pH 7.5, 500mM NaCl, 1mM DTT, 1% (v/v) glycerol. Fractions were assessed by SDS-PAGE and negative stain EM for the presence of complete DARP14-3G124Mut5 cages. Bound sfGFP V206A was evident by the green color and the occupancy was estimated by amino acid analysis performed at the UC Davis Molecular Structure Facility based on a least-squares fit to amino acid abundances.

### Negative stain EM

5 *u*L of fresh Superose 6 Increase eluent was adjusted to ~ 50 *u*g/mL, applied to glow-discharged 300 mesh formvar-carbon copper grids (Electron Microscopy Sciences) for one minute and blotted away. After two washes with filtered water, the grid was stained with 2% uranyl acetate for 30 sec. Images were taken on a Tecnai T12 or a TF20.

### Cryo-EM data collection

Fresh fractions from the Superose 6 Increase column containing DARP14-3G124Mut5 bound with sfGFP V206A were pooled and concentrated to 2 mg/mL. The sample was diluted to 1 mg/mL and final buffer composition of 10mM Tris pH 7.5, 500mM NaCl, 1mM DTT, 0.5% glycerol immediately prior to freezing. Quantifoil 200 mesh 1.2/1.3 copper grids (Electron Microscopy Sciences) was treated with 0.1% poly-lysine (Sigma-Aldrich) for 4-6 hours prior to freezing and cleaned of excess poly-lysine by washing with filtered water three times. 2.5 *u*L of sample was applied to the grids without glow discharging and frozen using a Vitrobot Mark IV (FEI). 1,929 movies were collected on a FEI titan Krios microscope (Thermo Fisher) with a Gatan K2 Summit direct electron detector in counting mode with image shift at a pixel size of 1.07 Å, and defocus values around -2.5 μm. Movies with 40 frames were collected over 8 sec with ˜7.00 e^-^*A^-2^*sec^-1^ dose rate.

### Cryo-EM data processing and model building

Raw movies were corrected for beam-induced motion using MotionCor2 (31) and the CTF estimation was performed with CTFFIND4 on non-dose weighted micrographs (32). 2D class averages of manually picked particles were used as templates in auto-picking in RELION 2.0 (24). Particles extracted from motion corrected, non-dose weighted micrographs that included frames 3-20 were passed into cryoSPARC (23). Two initial T symmetry enforced models were calculated *de novo* using *ab initio* reconstruction with a subset of total particles showing good 2D class averages. The good class from this reconstruction was selected and fed into a homogeneous refinement in cryoSPARC with a mask around the T33-21 core. These particles went through another round of 3D refinement for CTF and beamtilt refinement in RELION 3.0. Using chimera, densities corresponding to 11 DARPins and 11 sfGFP V206A were removed from the refined map to generate an asymmetric map that contained only the T33-21 core and one DARPin with a bound GFP. This asymmetric map was used to perform signal subtraction on T symmetry expanded particles (27) as previously described (26). 3D classification without alignment was performed on the post-signal subtraction particles. Two additional rounds of refinement with local angular searches were performed on the 3D class with the best density for the GFP in RELION 3.0. The first round of refinement used a mask that contained the T33-21 core and one DARPin with a bound GFP. The second round used a mask containing only one trimer of each type adjacent to the DARPin and GFP. The refined particles were subjected to a final round of multi-body refinement as previously described (33). We defined two bodies, one being the T33-21 core and the other being the density enclosed by the mask used in the second round of refinement. The data processing procedure is outlined in Fig. S2. Local resolution of the final map was established using the Resmap program (28) with resolution values sampled in increments of 0.25 Å.

All coordinate refinements were performed with phenix.real_space_refine (34). For the T33-21 core, the atomic coordinates of our previously determined T33-21 cryo-EM structure (PDB 6C9I) (10) were fitted into the cryoSPARC homogeneous refinement map in Chimera (35) and refined against this map. Additional residues in the DARPin region were built based on the previously determined DARP14 structure (PDB 6C9K) (10). For the model including the whole DARPin sequence, the body 1 map from multi-body refinement in RELION was sharpened with phenix.auto_sharpen (34) with resolution-dependent local sharpening and used for model building. The residues in 6C9K were corrected to match those in 3G124 and the added Mut5 mutations. The model was then trimmed to contain one trimer of subunit A, one trimer of subunit B, and one DARPin. The trimmed model was fitted into the sharpened map and refined iteratively with secondary structure restraints while allowing morphing or refinement at individual residues. Output from phenix was inspected manually and edited in COOT (36). Next, to properly fit a sfGFP structure (PDB 4W6B) (37) into the GFP density of the same map, we utilized an existing crystal structure –a complex between DARPin and eGFP (PDB 5MA8) (19). After aligning DARPin to DARPin in the DARP14-3G124Mut5 and 5MA8 structures, and replacing the eGFP in 5MA8 with the sfGFP in 4W6B, the sfGFP model was further refined with rigid body movement.

## SUPPLEMENTARY FIGURES

**Figure S1.**
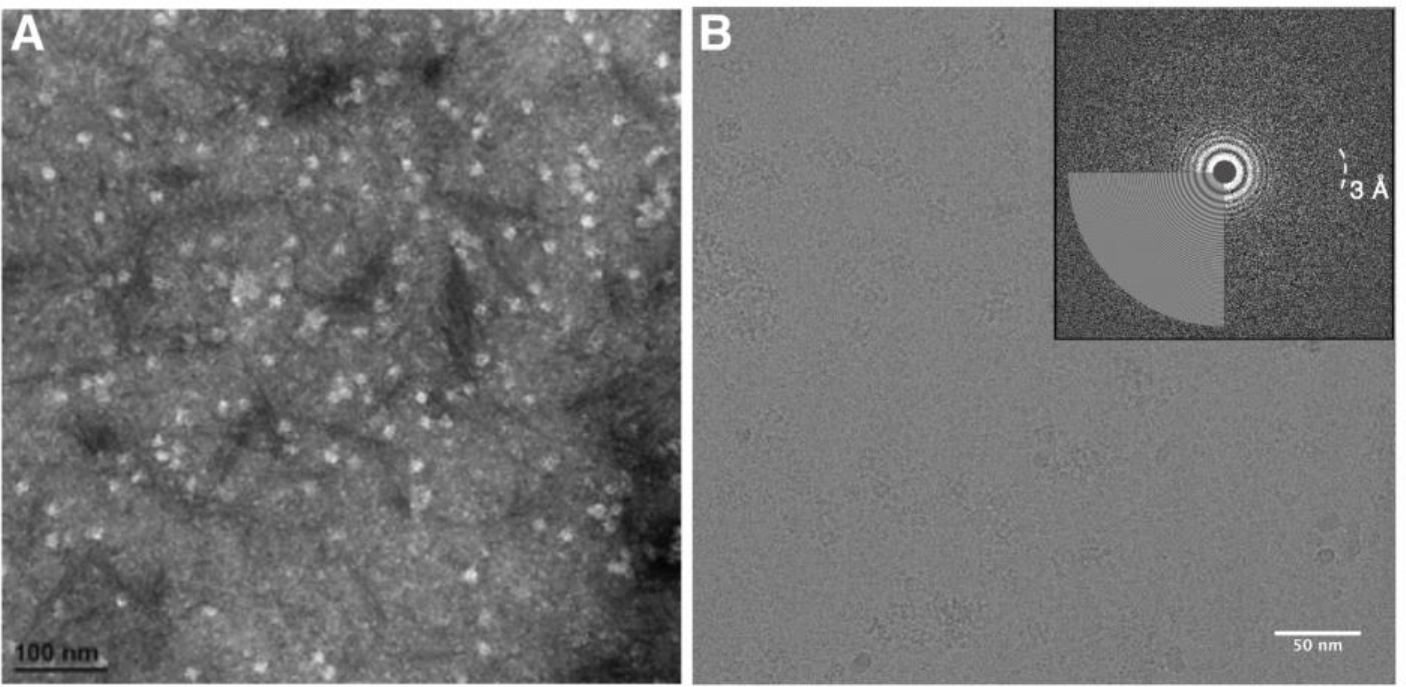
Preliminary biochemical analysis of the scaffold with its bound GFP. (A) Negative stain image of the complex. (B) Raw cryo-EM image with inset CTF correction result.

**Figure S2.**
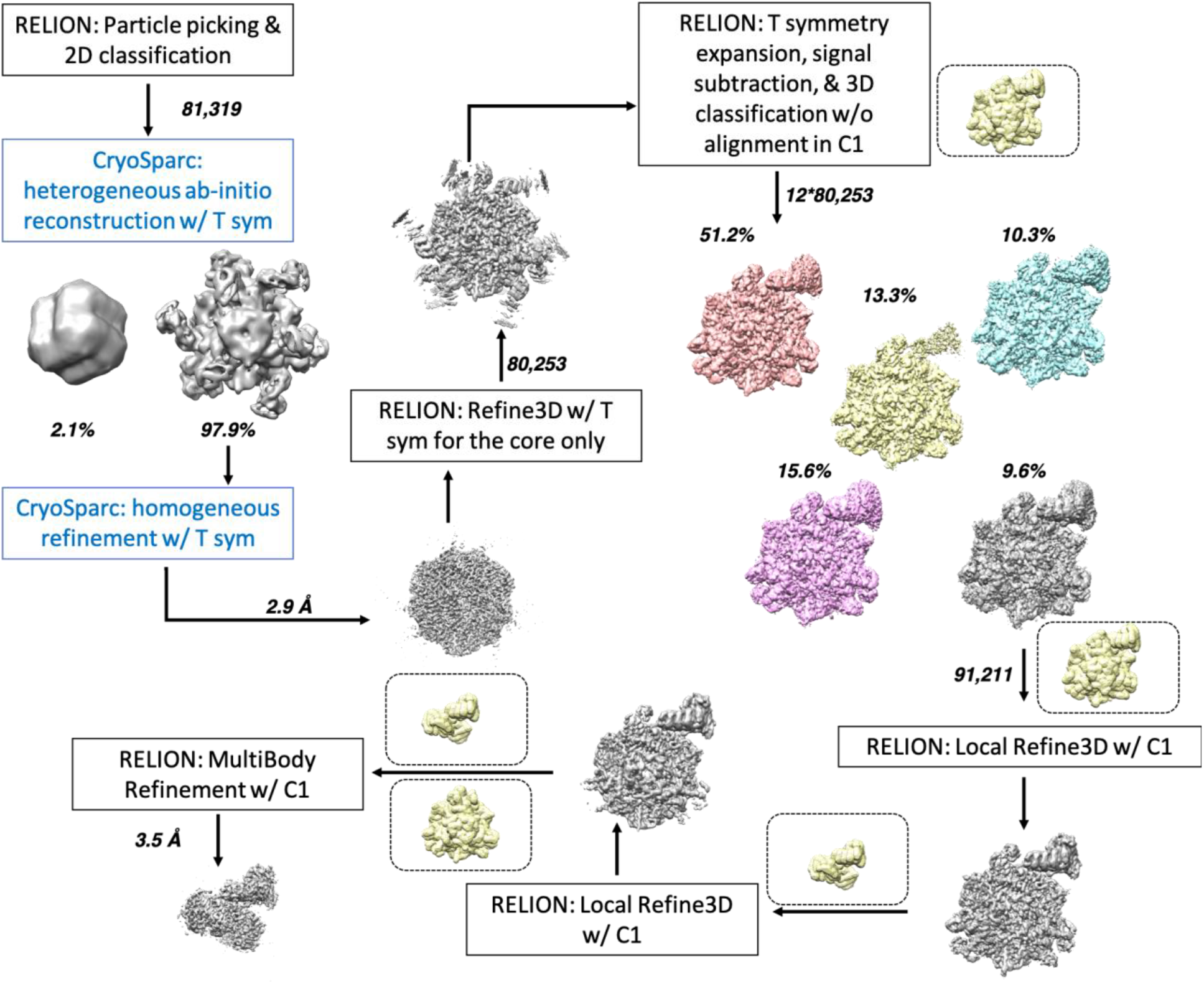
Cryo-EM data processing flow chart. Yellow densities inside dotted boxes indicate the masks used. For the 3D classification result, the coloring scheme is: class 1, grey; class 2, yellow; class 3, cyan; class 4, purple; class 5, pink.

**Figure S3.**
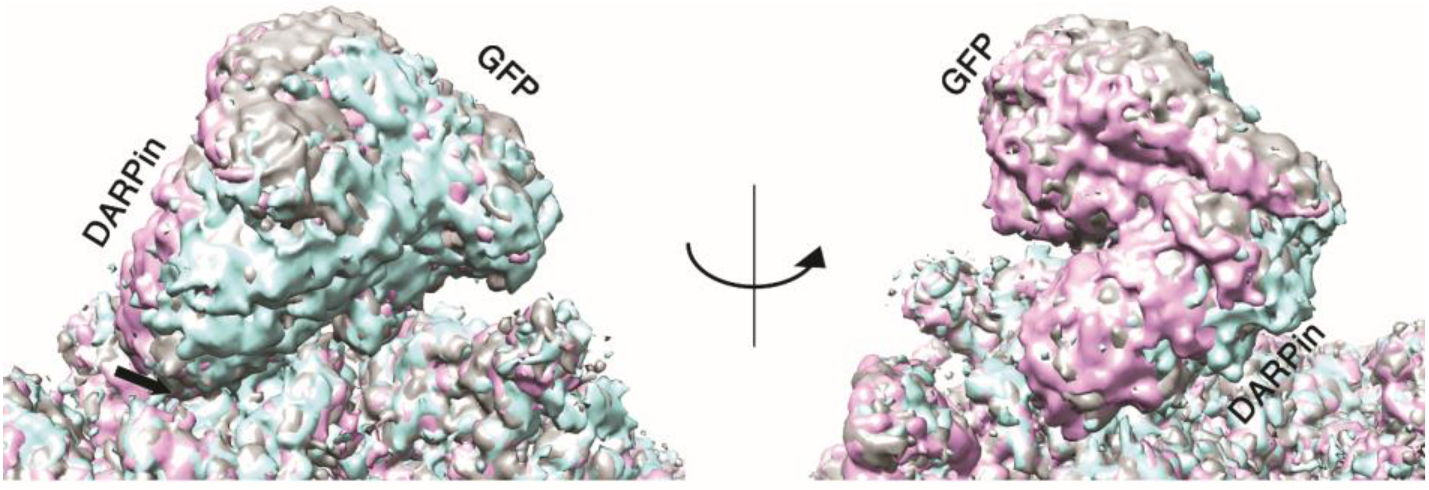
Comparisons between 3D classes (class 1, 3, and 5) of signal subtracted particles. Only a portion of the density is shown for clarity. An arrow highlights the secondary contact site between the DARPin (Gly 187) and Gly 108-Thr 109 on subunit A of the core assembly. Each class is colored the same as in Fig. S2.

## Supplementary text

### Protein sequences

- DARP14-3G124Mut5
  Subunit A: ~~~
MRITTKVGDKGSTRLFGGEEVWKDSPIIEANGTLDELTSFIGEAKHYVDEEMKGILEEIQNDIYKIMGEIGSKG
KIEGISEERIAWLLKLILRYMEMVNLKSFVLPGGTLESAKLDVCRTIARRALRKVLTVTREFGIGAEAAAYLLALS
DLLFLLARVIEIEQGKKLLEAARAGQDDEVRILMANGADVNAADDVGVTPLHLAAQRGHLEIVEVLLKCGAD
VNAADLWGQTPLHLAATAGHLEIVEVLLKNGADVNARDNIGHTPLHLAAWAGHLEIVEVLLKYGADVNAQ
DKFGKTPFDLAIDNGNEDIAEVLQKAA
~~~
  Subunit B: ~~~
MPHLVIEATANLRLETSPGELLEQANKALFASGQFGEADIKSRFVTLEAYRQGTAAVERAYLHACLSILDGRDI
ATRTLLGASLCAVLAEAVAGGGEEGVQVSVEVREMERLSYAKRVVARQRLEHHHHHH
~~~

- Super folder GFP V206A
  ~~~
MSKGEELFTGVVPILVELDGDVNGHKFSVRGEGEGDATNGKLTLKFICTTGKLPVPWPTLVTTLTYGVQCFSR
YPDHMKRHDFFKSAMPEGYVQERTISFKDDGTYKTRAEVKFEGDTLVNRIELKGIDFKEDGNILGHKLEYNFN
SHNVYITADKQKNGIKANFKIRHNVEDGSVQLADHYQQNTPIGDGPVLLPDNHYLSTQSALSKDPNEKRDH
MVLLEFVTAAGITHHHHHH
~~~

## Supplementary Tables

**Table S1.** CryoEM data table.

| Data collection/processing |  |  |
| --- | --- | --- |
| Microscope | Titan Krios |  |
| Voltage | 300 kV |  |
| Camera | Gatan K2 direct electron detector |  |
| Camera mode | Counting |  |
| Target defocus, $\mu\text{m}$ | -2.5 | |
| Exposure time per video, s | 8 |  |
| Total Dosage, $\text{e}^-/\text{\AA}^2$ | 56 | |
| Pixel Size, $\text{\AA}/\text{Pixel}$ | 1.07 | |
|  | <u>Cage core</u> | <u>Partial core, DARPIn, GFP</u> |
| Reconstructions |  |  |
| Software | RELION-2.0, cryoSPARC | RELION-2.0, RELION-3.0 |
| Symmetry | T | C1 |
| Particles | 80,253 | 91,211 |
| Overall resolution (masked), $\text{\AA}$ | 2.9 | 3.5 |
| Coordinate Refinement |  |  |
| Protein residues | 3432 | 1166 |
| Ramachandran, % |  |  |
| Favored | 96.5% | 95.3% |
| Allowed | 3.5% | 4.7% |
| Outliers | 0% | 0% |
| Model-map CC (masked) | 0.848 | 0.714 |

